# Molecular determinants of species-specific cell-cell recognition activating the class II gamete fusogen HAP2

**DOI:** 10.1101/2025.05.09.653094

**Authors:** Jennifer F. Pinello, Andrés Ferriño, Hibba Hussain, Pierre Legrand, Sujan Manikumar, Ruhama Demissie, Félix A. Rey, William J. Snell, Eduard Baquero

## Abstract

Species-specific adhesion of complementary gametes is a prerequisite to cell fusion during fertilization. Despite its importance, the molecular mechanism underlying cell–cell recognition and membrane adhesion to drive gamete fusion is not fully understood for any organism. In the alga, *Chlamydomonas*, the species-specific gamete adhesion protein MAR1 guides the fusogen HAP2 into a pre-fusion conformation on mt− gametes and serves as a receptor for the FUS1 adhesion protein on mt+ gametes. Here, we show that soluble recombinant MAR1 completely blocks the HAP2-dependent gamete fusion process by binding to FUS1 on mt+ gametes and report the X-ray structure of the MAR1-FUS1 complex. In vivo gamete fusion experiments showed that mutations of key residues at the observed MAR1-FUS1 interface strongly impair adhesion, yet gamete fusion still proceeds, highlighting the efficiency of the fusion-activation process. These findings uncover the molecular architecture of a surface receptor complex whose formation releases the viral class II viral fusion-protein ortholog HAP2, allowing it to undergo its membrane-fusogenic conformational change.

## INTRODUCTION

Fusion of haploid gametes to form a diploid zygote arose early in the evolution of eukaryotes. The membrane fusion reaction depends on the seamless integration of two functions -- species-specific attachment of the membranes and fusion of their lipid bilayers. Despite over 100 years of research on fertilization^1^, the mechanisms underlying these concerted adhesion and fusion events remain poorly understood^2^. In the sperm of both fish and mammals, the membrane recognition and adhesion apparatus is a conserved trimeric complex composed of membrane proteins TMEM81, SPACA6, and Izumo1^3–5^. Divergent egg proteins interact with this sperm protein complex – JUNO in mice and humans and Bouncer in zebrafish and medaka^3,6–10^. Although many other proteins are now known to be important for fertilization in chordates^11–20^, a full understanding of the gamete fusion reaction in these species has been hampered because the ultimate actor in the fusion reaction, the membrane fusogen, is unknown.

The only organism for which the essential elements of the gamete recognition, adhesion, and membrane fusion machinery have been identified is the unicellular green alga *Chlamydomonas reinhardtii* (hereafter, *Chlamydomonas*)^21–24^. When *Chlamydomonas* mt− and mt+ gametes are mixed to initiate fertilization, they interact through ciliary adhesion, thereby activating release of their cell walls and extension of membrane-enclosed, tubular projections – the mt+ and mt− mating structures – from the plasma membranes between the cilia^25^. The tip of the mt+ mating structure carries the mt+ adhesion protein, FUS1^23^, while the tip of the mt− mating structure contains the mt− adhesion protein MAR1^21^. MAR1 is bifunctional and is also an obligatory chaperone for the ancestral gamete fusogen, HAP2, on mt− gametes^21^. The continued motility of the cilia-tethered gametes thrusts their mating structures into contact^26^. Fusion between the tips of the mating structures is followed almost immediately by complete cellular coalescence to form the diploid zygote. Genetic disruption of the FUS1 gene in mt+ gametes or of the MAR1 gene in mt− gametes prevents mating structure adhesion and gamete fusion^21,23,24,26^. Disruption of HAP2 in mt− gametes has no effect on membrane adhesion but abrogates fusion^21,22,26^.

FUS1 is an ortholog of plant sperm GEX2 proteins, whose family members are found throughout the *Viridiplantae*^21,27–29^. Disruption of GEX2 in *Arabidopsis thaliana* strongly impairs sperm-egg fusion and seed set^29^. Contrary to the FUS1/MAR1 receptor pair in *Chlamydomonas*, no binding partner for a GEX2 in multicellular plants has been identified. FUS1/GEX2 family members are single trans-membrane domain proteins with a short cytosolic tail and a long ectodomain predicted to be composed of 7 tandemly arranged, Ig-like (IgL) domains (Fig. 1A)^21^. Unlike FUS1, MAR1 is a protein specific to green alga. It has a long, intrinsically disordered cytosolic tail and an ectodomain divided roughly into two halves, an N-terminal, membrane-distal, cysteine-rich region (CRR) and a membrane-proximal, extended proline-rich region (PRR, Fig. 1A).

**Figure 1.**
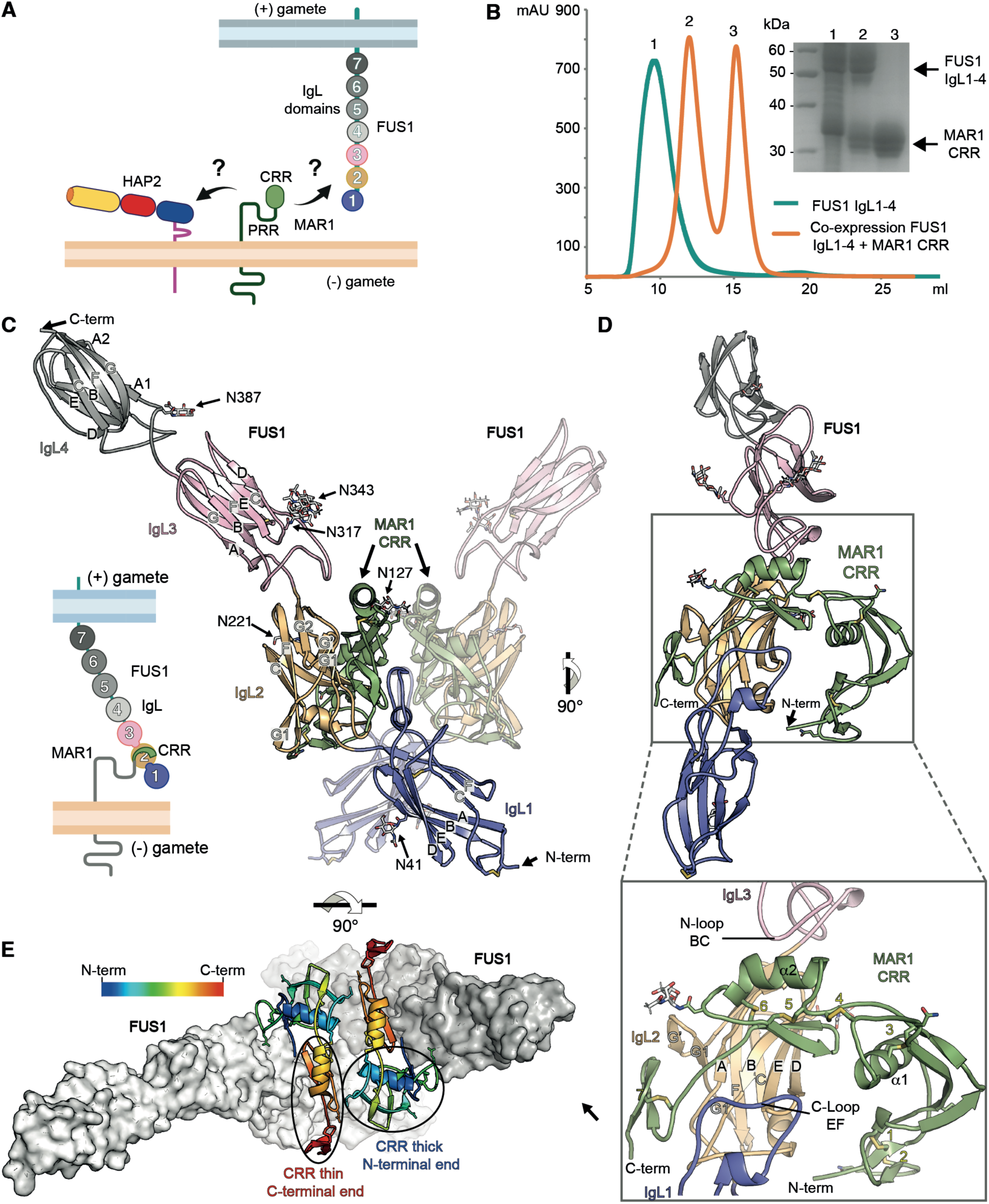
MAR1 CRR form a stable complex with FUS1 IgL1-3. (A) schematic representation of HAP2, MAR1 and FUS1, illustrating their relative positions on the membranes of *Chlamydomonas* mt− and mt+ gametes and the interactions of MAR1-HAP2 and MAR1-FUS1 (black arrows). (B) Size-exclusion chromatography profile of the complex purified from the supernatant of insect cells co-transfected with MAR1 and FUS1 constructs. The inset (right) shows the SDS-PAGE analysis of the main chromatographic peaks indicated in the elution profile (left). (C) Ribbon representation of the crystal structure of the MAR1 CRR-FUS1 IgL1-4 complex. The CRR is colored green and the IgL domains (1-4) are colored blue, yellow, pink and gray, respectively. One CRR-FUS1 IgL1-4 pair from the asymmetric unit is highlighted. Visible β-strands of each IgL are labeled with capital letters. N-glycans are shown in stick representation and labeled according to the corresponding asparagine residue to which they are linked. (D) The highlighted pair CRR-IgL1-4 from panel C is shown rotated by 90°, revealing CRR secondary structure. The inset provides a zoomed-in view of the complex highlighting the secondary structure elements involved in the interaction. E) Top view of the complex from panel C, with CRR molecules colored in rainbow gradient from N-terminal (blue) to C-terminal (red). FUS1 molecules are shown in white surface representation.

HAP2 (also termed GCS1^30^); is an ancient, conserved single-pass transmembrane protein with orthologs in archaea^58^ and among the deep-rooted branches of the eukaryotic tree. Some sub-branches appear to have lost or exchanged the gene. For example, among the Amorphea branch, HAP2 is found in amoeba and arthropods, but not in nematodes, fungi, or chordates, and a bona fide gamete fusogen in these organisms remains unidentified^31–35^. The protein was first identified in *A. thaliana* in a screen for male sterile mutants and shown to be important at a late step in fertilization^30,36^. Studies on fertilization in *Chlamydomonas* and in the murine malaria organism, *Plasmodium berghei*, demonstrated that disruption of HAP2 had no effect on gamete adhesion, but abrogated membrane bilayer merger and zygote formation^22,34^. Subsequent work has shown that HAP2 is expressed in gametes of the cnidarian, *Hydra*^37^, and the apicomplexan, *Cryptosporidium parvum*^38^ and is essential for fertilization in *Nematostella vectensis*^39^, *Babesia bovis*^40–43^, *Tetrahymena thermophila*^44–47^, *Dictyostelium discoideum*^48^, and *Drosophila melanogaster*^49^. Antibodies against HAP2 of the malaria parasite genus, *Plasmodium*, potently reduce mosquito transmission of malaria^50–52^, establishing HAP2 as a candidate for a transmission-blocking malaria vaccine.

Notably, HAP2 is a structural homolog of the class II membrane fusion proteins (MFPs)^32,53,54^ including proteins within the cellular family of FF fusogens^55–57^, and those proteins coating the envelopes of certain viruses, including yellow fever^59,60^, rubella^61,62^, and chikungunya^63^ viruses. The viral class II MFPs heterodimerize co-translationally with a chaperone accompanying protein (AP) in the viral envelope^64^. Upon virus interactions with a target cell receptor and endocytosis, the acidic endosomal environment induces MFP/AP heterodimer dissociation. Released from the AP chaperone’s grip, the MFP’s fusion loop inserts into the endosomal membrane, adopting a transient, homotrimeric elongated conformation and bridging the 10-15 nm gap between the two membranes^65,66^. The trimer then undergoes a structural transition into its lowest energy conformation by folding its three protomers into a “hairpin” that forces the two membranes against each other leading to the formation of a fusion pore^67^ and cytoplasmic continuity. Previous studies reported that adhesion between mt− and mt+ gametes was sufficient to induce HAP2 trimerization and membrane fusion, however the molecular details of this triggering are unknown^68^.

As with the viral MFPs, HAP2 is expected to drive gamete fusion by undergoing a major conformational change from a metastable form into a stable, post-fusion trimer of hairpins. The reported HAP2 structures mainly show the typical “post-fusion” trimer of hairpin conformation^69–72^, yet a structure of a monomeric, pre-fusion form of HAP2 was also reported^73^. One fundamental distinction from its viral counterparts is that the HAP2 fusogenic conformational change is triggered by contact between the plasma membranes of the gametes and not by the low pH of the endosomal compartment. Although the nature of the MAR1-HAP2 interaction has not been characterized, the MAR1-FUS1 interaction was shown to be direct, and recombinantly expressed ectodomains of MAR1 and FUS1 bind to each other in vitro^21^. Thus, characterizing the MAR1-FUS1 recognition mode at the center of this concerted membrane adhesion and bilayer merger reaction is central to understanding regulation of HAP2-driven membrane fusion.

Here, using biochemistry, X-ray crystallography and mutagenesis in vitro and in vivo experiments we identify the minimal interacting regions of MAR1 and FUS1 and report the structural basis of their interaction. Our discovery that HAP2-driven fusion is resistant to adhesion-impairing MAR1 mutations suggests a differential requirement for HAP2 activation and efficient cell adhesion, as only a few activated HAP2 molecules would be required for gamete fusion to proceed. The structure of the MAR1-FUS1 receptor complex provides a unique entry point for understanding the mechanisms that couple gamete membrane adhesion interactions to the triggering of the HAP2 membrane-fusogenic conformational change, a step not elucidated for any non-viral organism to date.

## RESULTS

### Structure determination

We engineered multiple constructs for the heterologous expression and secretion of the MAR1 and FUS1 ectodomains into the supernatant of transfected eukaryotic cells (See Methods). Both ectodomains exhibited high secretion yields, although FUS1 displayed a strong tendency to aggregate. Co-transfection with the MAR1 construct prevented FUS1 aggregation thanks to their interaction. Co-transfection with just the MAR1 CRR construct (residues 26-161; numbering includes the signal sequence) also avoided aggregation of the FUS1 ectodomain (IgL1-7, residues 17-775) or a truncated IgL1-4 (residues 17-479) version. The strep-tagged MAR1 CRR purified by tag-affinity chromatography co-eluted with 6xHis-tagged FUS1. Subsequent size exclusion chromatography showed two elution peaks, one corresponding to the complex and one to excess MAR1 (Figure 1B). In sum, co-expressing the MAR1 CRR with FUS1 IgL1-4 resulted in a purified complex, indicating that their direct interaction takes place via the N-terminal, membrane distal halves of the two proteins. We obtained crystals of the MAR1 CRR / FUS1 IgL1-4 complex diffracting to 3.2 Å resolution, which allowed the structure determination by molecular replacement. The diffraction data collection, structure determination and model refinement statistics are listed in Table S1.

### FUS1 organization

The structure showed that each FUS1 IgL domain adopts a *β*-sandwich fold with a 4-stranded “ABED” sheet opposing a 3-stranded “CFG” sheet (see Figure 1C and S1A). This organization is related to “set C1” of constant domain immunoglobulins^74,75^, though it lacks the conserved inter-sheet disulfide bond connecting strands B and F. Instead, FUS1 IgL1 contains two disulfide bonds at non-standard locations that are conserved among FUS1 proteins in other Chlorophytes, whereas the other IgLs lack disulfide bonds (Figure S2). IgL1 also lacks the C-terminal *β*-strand G, where the polypeptide runs as random coil roughly maintained at the corresponding location by the short *β*-strand G′ interacting with *β*-strand A. Each IgL domain features three “N-loops” (connecting *β*-strands B to C, D to E and F to G) and three “C-loops” (connecting *β*-strands A to B, C to D and E to F), named for their proximity to the domain’s N- or C-termini, respectively. The linker between domains, as well as the N- and C-loops are substantially elaborated, displaying additional secondary structure elements (Figure 1C). In the FUS1 rod-like molecule, IgL1 exposes its N-loops at the membrane-distal end (note that in the related immunoglobulins, the N-loops of the variable domains carry the antigen complementarity determining regions, or CDRs), while its C-loops interact with the N-loops of IgL2. Similarly, the IgL2 C-loops engage with the N-loops of IgL3 making a rigid IgL1-3 rod through stable interdomain interactions. At the interface between IgL3 and IgL4, the interacting C- and N-loops are shorter, resulting in a looser contact that increases the mobility of IgL4. Within this domain arrangement, IgL1, 2 and 3 orient their ABED in the same direction, while the loosely linked IgL4 is rotated 180 degrees.

Using the FUS1 structure, Foldseek^76^ identified homology to the plant GEX2s discussed above^21^, displaying E-values < 10e-21 and amino acid sequence identity of ~14-16%). Structural comparison of FUS1 with AlphaFold-predicted models^77^ of GEX2 proteins revealed that FUS1 IgL 1-3 align with consecutive GEX2 IgL domains of several species including *Panicum hallii* (IgL 1-3), *Zea mays* (IgL 3-5), *A. thaliana* (IgL 2-4), *Capsicum baccatum* (IgL 3-5) and *Citrus clementina* (IgL 3-5) (Figure S2). These domains share conserved sequence motifs, indicating a common evolutionary origin (Figure S2). GEX2 proteins do not display the two conserved disulfide bonds in the equivalent FUS1 IgL1, instead most GEX2 proteins display a single cysteine in the vicinity of the *β*-strand A, near the position of the first cysteine of FUS1 disulfide bond 1 (Figure S2). Other GEX2 domains also display free cysteines and with exception of *P. hallii* GEX2, there is a conserved pair of cysteines in the equivalent of FUS1 IgL3 between *β*-strands F and G, which are predicted by AlphaFold3 to form a disulfide bond. Foldseek also identified FUS1 homology to the IgL domains of actin-binding filamins B and C (E-values < 1- e-17 and 15% aa seq. identity). Consistent with their cytosolic location, and contrasting with the FUS1 and GEX2 predictions, the filamin IgLs lack disulfide bonds, their N- and C-loops are shorter, and their interdomain connections are highly flexible.

### MAR1 organization

The structure revealed that the MAR1 CRR is a hook-shaped molecule with an overall novel fold stabilized by seven disulfide bonds. It contains two *α*-helices and four 2-stranded *β*-sheets, with the topology illustrated in Figure S1A. The molecule is asymmetric with an N-terminal thick end and a thin C-terminal end (the “hook tip”) (Figure 1D and 1E). Despite its overall novel fold, a Foldseek search identified weak similarity of the thin, C-terminal end (residues 109-162) with the epidermal growth factor-like (EGF-like) repeats of the extracellular matrix protein Fibulin-1 (E-value ~10e-1; 22-26% aa seq. id.) (see Figure S3A). Fibulin-1 is an extracellular matrix protein with nine tandemly arranged EGF-like repeats^78^ each containing 3 disulfide bonds, which correspond to disulfides 5 to 7 of MAR1, but lacking the MAR1 *α*-helix (helix *α*2) connecting the first two disulfides of the motif (Figure 1D and S3A).

### The MAR1/FUS1 complex

The structure showed that the MAR1/FUS1 complex is organized around a central antiparallel MAR1 CRR dimer, where the CRR protomers interact laterally through their central part (Figure 1C and 1E). In this arrangement, the thicker N-terminal end of one hook-like CRR protomer pairs with the thinner hook tip of the other, exposing both concave surfaces on the same side. The MAR1 hook tip inserts into a FUS1 groove whose floor is the exposed face of the ABED *β*-sheet of IgL2, and whose walls are formed by IgL1 C-loop EF and IgL3 N-loop BC – both of which are long and elaborated (Figures 1D and S1A). The thick end of MAR1 interacts with the IgL1 C-loop EF of the other FUS1 protomer, in such a way that each MAR1 molecule interacts with both FUS1 protomers (Figure 1E). The total buried surface area in the complex is 1343 Å^2^ for FUS1 and 1218 Å^2^ for MAR1 and involves 10 hydrogen bonds between side chains of FUS1 residues mostly interacting with the main chain of the MAR1 hook tip (Figure S1B and S3B). A key feature of the interaction is the proline ring of P142 inserted snuggly into a cage of FUS1 aromatic side chains on the groove floor, which also interacts with the hydrophobic side chain of MAR1 M141 (Figure 2A). These interactions provide high specificity to the recognition between MAR1 and FUS1.

**Figure 2.**
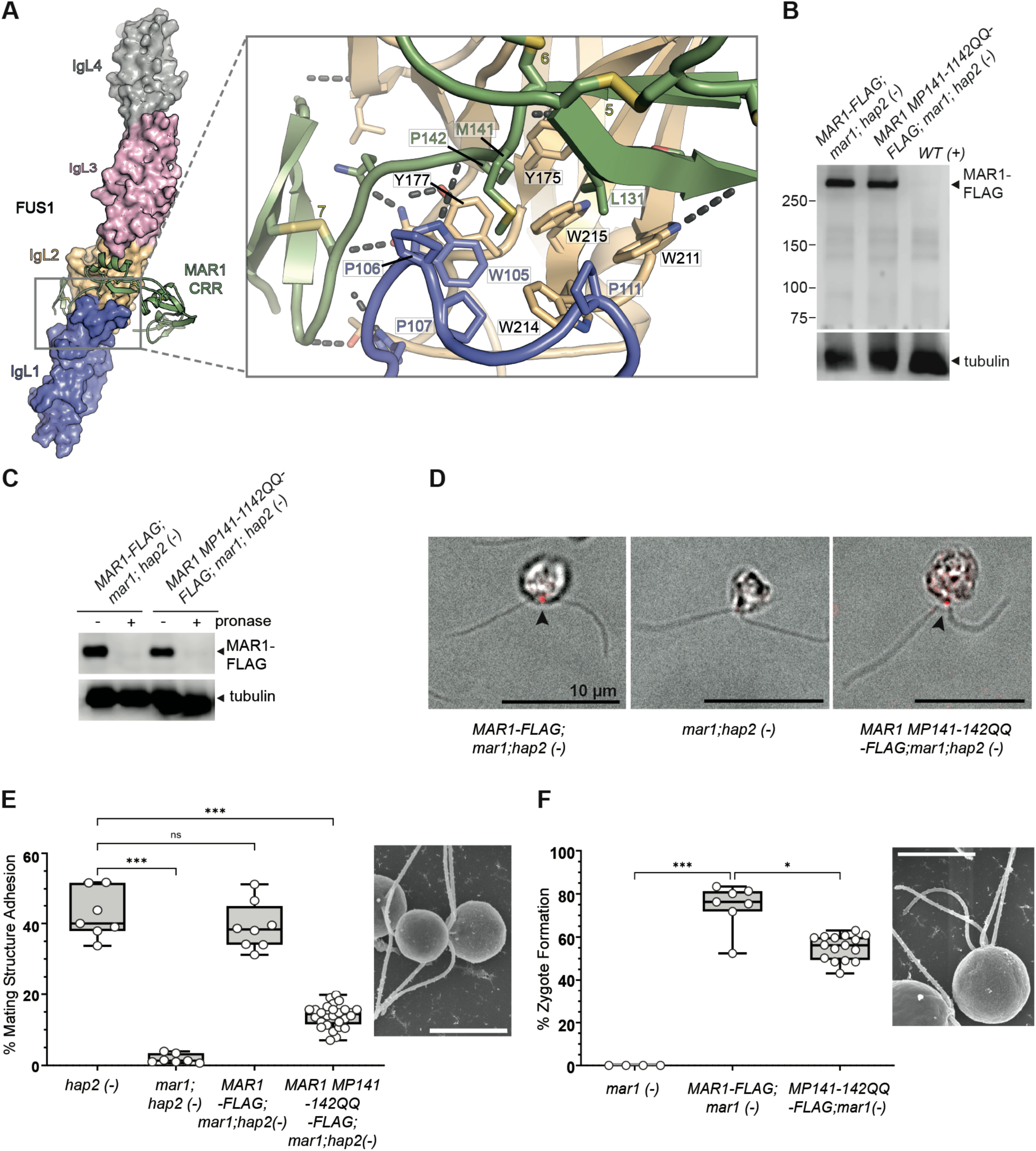
MAR1 MP141-142QQ mutations disrupt *Chlamydomonas* mating structure adhesion and gamete fusion. (A) The CRR-IgL1-3 complex is shown in ribbon and surface representations respectively, using the color scheme in Figure 1. The inset provides a close-up view of the aromatic cage in the FUS1 groove, where MAR1 residues M141 and P142 insert. The residues participating in this interaction are labeled and shown as sticks. The dashed lines in the inset indicate hydrogen bonds between the MAR1 and FUS1 chains, which are listed in Figure S1. (B-D) *MAR1 MP141-142QQ-FLAG*(-) gametes express MAR1 similarly to wild-type *MAR1-FLAG*(-) gametes. (B) Representative immunoblots of lysates of gametes expressing *MAR1-FLAG*, *MP141-142QQ-FLAG*, or no MAR1 protein. (C) The sensitivity of the *MAR1 MP141-142QQ-FLAG* protein to pronase treatment indicates that it was expressed at the cell surface of mt− gametes. (D) Representative composite confocal immunofluorescence and DIC images show localization of MAR1-FLAG protein (red) at the mating structures of wild-type and mutant *MAR1-FLAG(-)* gametes, but not in *mar1;hap2*(-) gametes. (E-F) Mating structure adhesion (E) and fusion (F) are reduced in *MAR1 MP141-142QQ-FLAG* mt− gametes mixed with WT mt+ gametes. Representative SEM images of an adhering pair of mt+ and mt− gametes and a zygote, respectively, are shown to the right of plots of pair formation (E) and zygote formation (F) of the indicated samples. Individual biological replicates (n) are shown as open circles. Statistical significance was assessed using a Kruskal-Wallis test with Dunn’s multiple comparisons test: *p < 0.05, ***p < 0.001; ns, not significant. Representative blots and images shown are the results of 2-4 independent experiments. Scale bars are 10 µm.

### mt− gametes expressing MAR1 protein with mutations in the binding interface with FUS1 are impaired in mating structure adhesion

To confirm the functional relevance of the MAR1 141-MP-142 insertion into the FUS1 aromatic pocket, we examined mating structure adhesion in samples of mt+ gametes mixed with *mar1*;*hap2* mt− gametes expressing either wild-type MAR1-FLAG or a MAR1-FLAG with two consecutive glutamines replacing the methionine-proline motif at MAR1 positions 141-142. The endogenous *MAR1* and *HAP2* genes in these mt− gametes had been disrupted^21,79,80^. The lack of HAP2 expression, which has no effect on MAR1 expression or localization^21^, rendered the gametes incapable of fusion, and therefore allowed assessment of mating structure adhesion alone. Immunoblot analyses of gametes expressing the wild-type and mutant proteins showed that MAR1 141-QQ-142-FLAG was present at the same levels as wild-type MAR1-FLAG (Figure 2B). In addition, like the wild-type protein, MAR1 141-QQ-142-FLAG was susceptible to degradation upon incubation of live gametes with pronase (Figure 2C), indicating that the protein was present at the cell surface. Immunostaining with anti-FLAG antibodies confirmed that MAR1 141-QQ-142-FLAG was correctly localized in between the bases of the 2 cilia, the site of the mt− mating structure (Figure 2D).

To assess mating structure adhesion, *mar1*;*hap2* mt− gametes expressing wild-type MAR1-FLAG or MAR1 141-QQ-142-FLAG were mixed with mt+ gametes and after 10 m, the reaction was stopped by addition of glutaraldehyde followed by quantification of the percent of cells adhering by their mating structures (Figure 2E). In the mixtures of wt+ gametes with *hap2* mt− gametes, over 40% of the gametes adhered by their mating structures (Figure 2E); whereas in the mixtures of mt+ gametes with *mar1*;*hap2* mt− gametes, mating structure adhesion failed to occur. Re-introduction of wild-type MAR1-FLAG into the *mar1*;*hap2* gametes rescued mating structure adhesion to control levels, but mating structure adhesion was reduced 3-fold compared to controls in the *mar1*;*hap2* gametes expressing MAR1 141-QQ-142-FLAG. Thus, the 141-MP142 residues have a critical function in the FUS1-MAR1 adhesive interaction. We next quantified the effects of mutation of MAR1 141MP142 to QQ on gamete fusion by expressing the proteins in *mar1* mt− gametes whose HAP2 protein was functional (Figure 2F). As expected from previous results, in mixtures of wt+ gametes with *mar1* mt− gametes, zygote formation failed to occur; whereas in the mixtures of mt+ gametes with *mar1* mt− gametes expressing wild-type MAR1-FLAG, ~75% of gametes formed zygotes. Interestingly, unlike the 3-fold reduction in adhesion in the MAR1 MP141-2QQ mutant gametes, we found fusion was reduced less than 30% in these mutants, indicating the plasticity and efficiency of the HAP2-mediated fusion reaction under conditions of suboptimal membrane adhesion.

### Incubation of mt+ gametes with wild type (WT), but not 141-QQ-142 mutant forms of the recombinant MAR1 CRR (rMAR1), blocks mating structure adhesion, HAP2 trimer formation, and gamete fusion

Immunofluorescence studies showed that rMAR1 (residues 26-161) bound to the mating structure of *WT* mt+ gametes, but not to the mating structure of *fus1* mt+ gametes, which lack the FUS1 protein (Figure 3A). As expected, rMAR1 had no effect on ciliary adhesion (Figure S4A), but almost completely blocked adhesion of the mating structures (Figure 3C, phase contrast images and Figure 3D, quantification). Microscopic assessment showed that the percentage of gametes undergoing mating structure adhesion was over 50% in buffer control samples without recombinant protein, and that the wild-type rMAR1 reduced mating structure adhesion 10-fold, to less than 5%. Further confirming the functional importance of residues at the FUS1-MAR1 interface seen in the X-ray structure, rMAR1-M141Q and rMAR1-P142Q mutants failed to block adhesion, whereas rMAR1-A111D, with a mutation in a residue distal to the interface, was fully capable of blocking adhesion of the mating structures (Figure 3C and 3D). Results from corresponding in vitro experiments mirrored the in vivo results. When WT or mutant forms of strep-tagged rMAR1 was pulled down from supernatants of *Drosophila* Schneider 2 cells co-expressing His-tagged rFUS1, immunoblotting of the precipitates with anti-His antibodies revealed that rFUS1 bound to WT rMAR1 and rMAR1-A111D, but failed to bind to rMAR1-M141Q and rMAR1-P142Q (Figure 3B).

**Figure 3.**
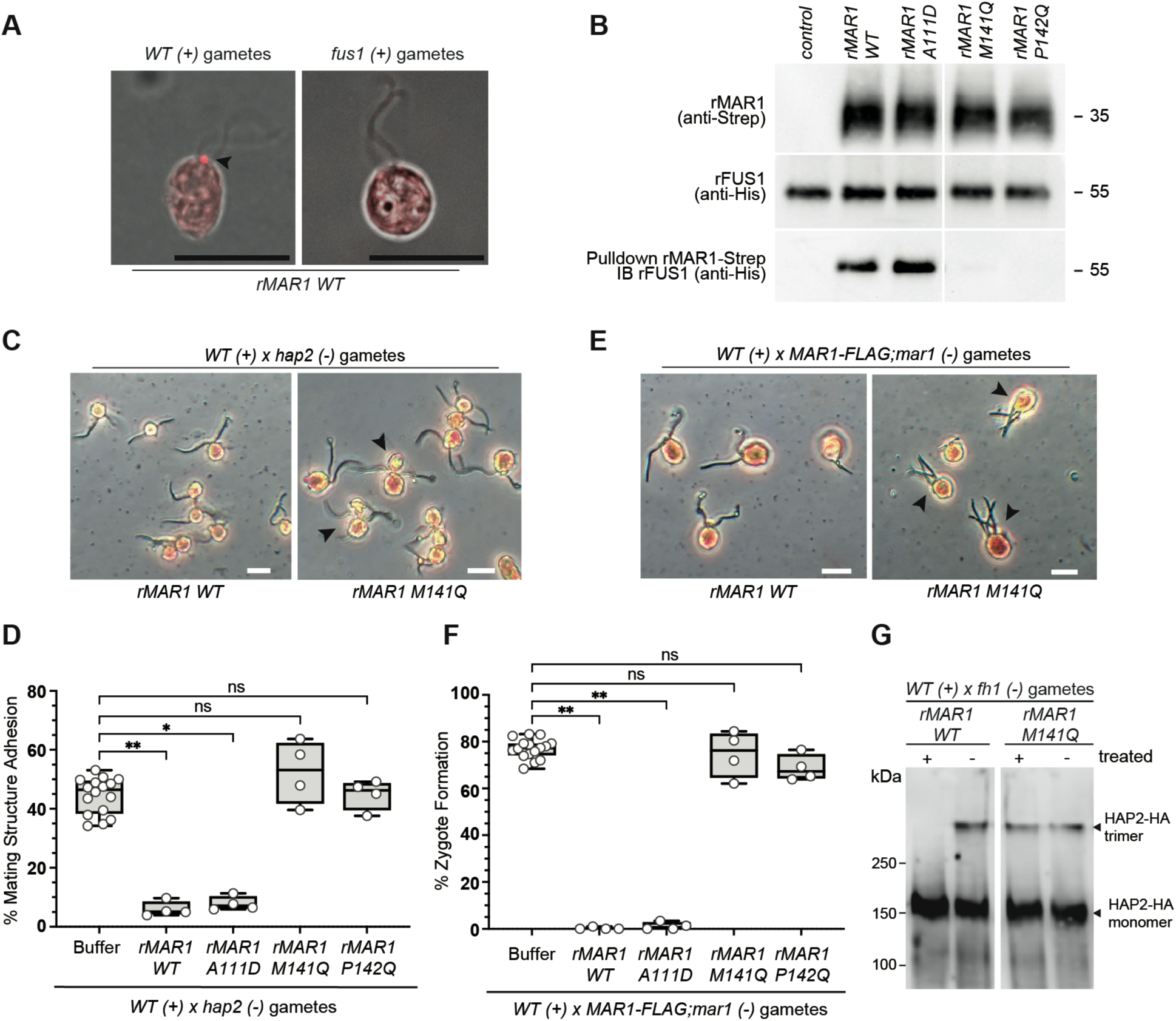
The recombinant MAR1 CRR (rMAR1) blocks adhesion of the mating structures, gamete fusion, and concomitantly, no HAP2 trimerization is observed. (A) rMAR1 binds to FUS1 on mt+ gametes. Representative composite confocal immunofluorescence and DIC images show rMAR1 bound to activated wild-type WT(+) but not *fus1(*+) gametes. Scale bars are 10 µM. (B) rFUS1 binds to WT rMAR1 but not to rMAR1-M141Q or -P142Q. Immunoblotting detected the presence/absence of rMAR1 (top) and rFUS1 (middle) in cell lysates. Eluates from rMAR1 pull down were immunoblotted (IB) with anti-HIS antibodies for detection of co-precipitating forms of rMAR1. (C-F) Gamete adhesion and fusion are blocked by WT rMAR1, but not by rMAR1 with single point mutations at the MAR1-FUS1 interface. After incubation with the indicated rMAR1 proteins with activated mt+ gametes, mt− gametes were added that were competent for mating structure adhesion only (*hap2(-)* gametes, C-D) or for adhesion and fusion (*MAR1-FLAG*;*mar1(-)*, E-F) and assessed for pair formation (C-D) or zygote formation (E-F). Representative phase contrast images of gametes (C, E; scale bars are 100µM). Box-and-whisker plot quantification (D,F) of rMAR1-treated or control cell mixtures with individual biological replicates (n) shown as open circles. Statistical significance was assessed using a Kruskal-Wallis test with Dunn’s multiple comparisons test: *p < 0.05, **p < 0.01, ns, not significant. (G) HAP2 trimer formation is blocked by pre-incubation with WT rMAR1 but not with interface mutant forms of rMAR1. Activated mt+ gametes co-incubated WT rMAR1, rMAR1-M141Q, or buffer were mixed with *fh1* mt− gametes^26,71^ followed by immunoblot analysis of cell lysates for HAP2 trimer formation. Representative blots and images shown are the results of 2-4 independent experiments. Scale bars are 10 µm.

Additional in vivo experiments in which mt+ gametes were mixed with fusion-competent WT mt− gametes instead of fusion-defective *hap2* mt− gametes demonstrated that the strong inhibition of mating structure adhesion by wild-type rMAR1 was accompanied by a nearly complete block to gamete fusion. 80% of cells in the control mixtures of gametes and in mixtures containing rMAR1 M141Q or rMAR1 P142Q fused to form zygotes, but less than 1% of the cells in samples containing WT rMAR1 or rMAR1-A111D formed zygotes (Figure 3E, phase contrast images and Figure 3F, quantification). Finally, blocking the FUS1-MAR1 interaction in these experiments with WT rMAR1 also blocked HAP2 trimer formation (Figure 3G), providing direct evidence that this specific FUS1-MAR1 interaction triggers the fusogenic conformational change of HAP2.

## DISCUSSION

Gamete membrane adhesion and lipid bilayer merger are seamlessly integrated during fertilization in unicellular and multicellular organisms. In vertebrates, adhesion complexes have been described but no bona fide gamete fusogen has been identified. In other taxa, HAP2 is the gamete fusogen but the molecular architecture of the receptor pair has been unknown. Our study reveals the structural and functional details of gamete adhesion recognition in *Chlamydomonas* providing a more complete understanding of a gamete fusion machinery. In this recognition, the tip of the MAR1 CRR hook inserts into a FUS1 groove in IgL2 flanked by loops projecting from IgL1 and IgL3. The interaction involves a network of hydrogen bonds established between the CRR main chain and FUS1 side chains together with the MAR1 motif 141-MP-142 inserting into a FUS1 aromatic pocket. This last interaction provides additional specificity to the mt− and mt+ recognition as the 141-QQ-142 mutation induced a 3-fold reduction in mating structure adhesion together with a 30% reduction in gamete fusion (Figure 2). These findings also demonstrate that impaired gamete adhesion remains sufficient for a HAP2-driven bilayer fusion reaction.

Fusion arrest was achieved by pretreating mt+ gametes with the recombinant MAR1 CRR domain. This blockade is the result of a complete occupation of all available FUS1 molecules by the rMAR1 CRR, precluding any interaction with MAR1 in the mt− gamete, demonstrating that gamete recognition, specifically mediated by the MAR1-FUS1 interaction, indeed activates the gamete fusion reaction. Consequently, FUS1 displaces HAP2 from its prefusion complex with MAR1, enabling it to insert the fusion loops into the mt+ membrane and undergo its energetically downhill conformational change to reach its final, lowest energy trimeric hairpin post-fusion conformation (Figure 4) that leads to membrane fusion. This process in *Chlamydomonas* bears similarities to the mechanisms of the structurally related viral class II MFPs where the MFP is maintained in a metastable pre-fusion conformation by interaction with an accompanying protein. Of note, paramyxoviruses, which fuse for entry at the plasma membrane and not in an endosome, have a structurally unrelated class I membrane fusion machinery in which the MFP relies on receptor binding by a viral protein. The two viral proteins are weakly associated at the viral surface, and this interaction maintains the MFP in a pre-fusion conformation until the companion protein recognizes a specific receptor at the surface of a susceptible cell^81,82^. Receptor binding by the paramyxovirus-accompanying protein triggers the conformational change of the MFP^83^, in a striking parallel to the *Chlamydomonas* HAP2/MAR1 pair and the recognition of FUS1 by MAR1.

**Figure 4.**
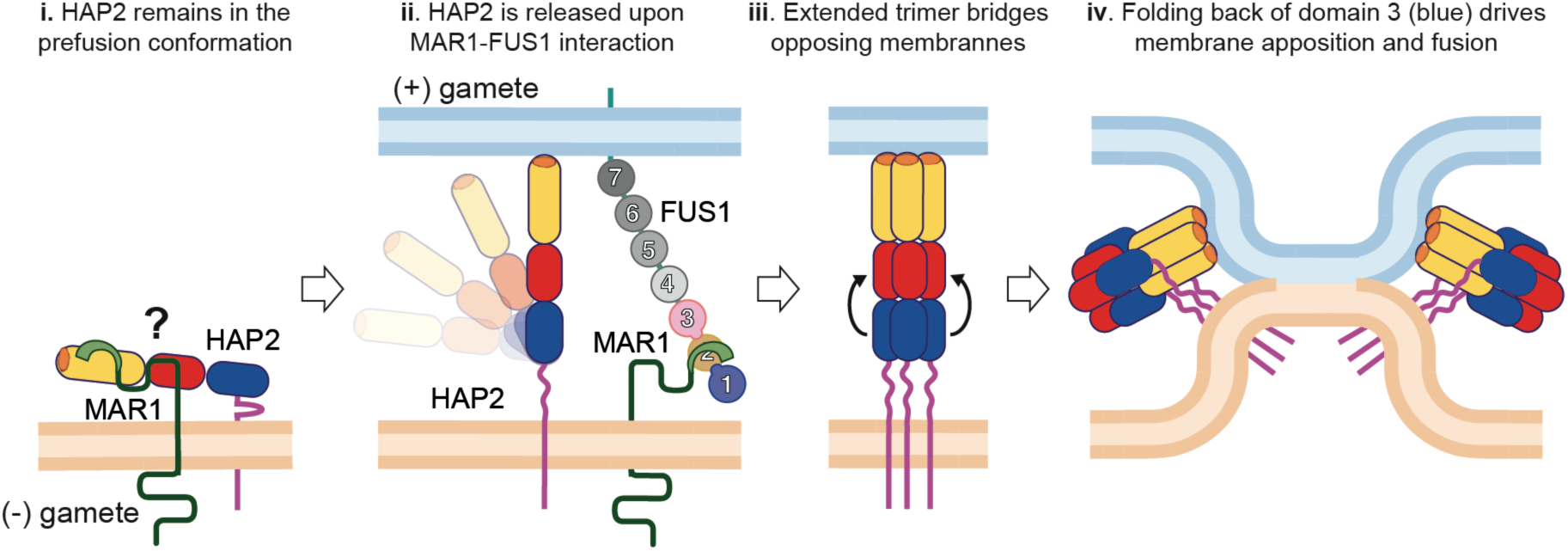
Proposed sequence of molecular events during *Chlamydomonas* gamete fusion. Cartoon based on the current understanding of virus fusion of paramyxovirus MFPs^92^ and class II virus MFPs^66,67^. Although a direct interaction between HAP2 and MAR1 has not be demonstrated, it was shown that HAP2 requires the presence of MAR1 to localize at the mating structure, and that both proteins can be co-precipitated under certain conditions^21^. We postulate that MAR1 accompanies HAP2 to the fusion site and interferes with fusion-loop exposure. i) Pre-fusion form. In this cartoon, HAP2 is represented as a rod-like molecule with its three main domains (I, II and III in red, yellow and blue, respectively, and the fusion loop in orange and the extensive “stem” region as a magenta line). MAR1 is drawn in green with an extended PRR hypothetically directing the CRR hook’s interference with HAP2 fusion loop exposure. The question mark indicates that the details of this interaction are unknown. II) Binding to FUS1 induces MAR1 to release HAP2, allowing fusion loop insertion into the opposite gamete, and thereby bridging the two membranes. II) A transient core extended trimer forms, bridging the two membranes, in which domain III (in blue) can relocate to a binding site at the domain I-domain II (red-yellow) interface. Curved arrows indicate the subsequent relocation of domain III to make a hairpin. III) harpin formation pulls the two membranes against each other, resulting in membrane apposition followed by their hemi-fusion (as indicated) and ending by formation of a fusion pore (not represented), in which HAP2 has reached its post-fusion, SDS-resistant trimer of hairpins (identified in the gel of Fig. 3G). The forces that drive the subsequent expansion of the initial fusion pore to whole cell fusion have not been so far explored.

Among the three elements of the *Chlamydomonas* fusion machinery, MAR1 is notably underrepresented across taxa probably due to an inherent diversity of HAP2-associated proteins, whereas orthologs of HAP2 and FUS1 are found in other eukaryotic lineages. Both FUS1/GEX2 and HAP2 proteins share a conserved structural scaffold within their respective families. Yet, they feature variable loops mediating the interactions, such as the N- and C-loops in IgL1 and IgL3 in FUS1 contributing to MAR1 binding shown here, or the HAP2 fusion loops responsible for membrane insertion^71,84,85^. An important difference, however, is that GEX2 and HAP2 are found together in the same gamete in plants (the sperm), whereas in algae they are in opposite gametes. It is known that GEX2 is important for gamete adhesion in plants, but the molecule it recognizes in the opposite gamete is unidentified, and what may couple adhesion to fusion in that case remains elusive.

In summary, our results reveal the molecular details of the interaction between two proteins required for the adhesion of complementary gametes that triggers the conserved protein HAP2 to drive membrane fusion in the model organism, *Chlamydomonas*. This system provides a paradigm for understanding the related molecular mechanisms of gamete fusion in all agriculturally important, beneficial and harmful plants and their insect pests, and in globally devastating unicellular pathogens, such as *Plasmodium, Trypanosoma, Leishmania, Babesia* and their insect vectors, which rely on HAP2 for transmission.

## RESOURCE AVAILABILITY

### Lead contact

Further information and requests for resources and reagents should be directed to and will be fulfilled by the lead contacts; Drs. William J. Snell and Eduard Baquero.

### Materials availability

Materials generated in this study including plasmids and *Chlamydomonas* strains are available upon request from the lead contacts or by direct purchase from the *Chlamydomonas* Resource center (https://www.chlamycollection.org/).

### Data and code availability

No new code or large-scale datasets were generated in this study. All data reported in this paper and any additional information required to reanalyze these data are available from the lead contacts upon request. Structural information has been submitted to the Protein Data Bank (PDB).

## EXPERIMENTAL MODEL AND SUBJECT DETAILS

### *Chlamydomonas* cells and cell culture

Cells were grown vegetatively in liquid TAP media at 22°C in a 13:11 light-dark cycle. Gametogenesis was induced by transfer of cells to nitrogen-free medium followed by overnight agitation on a shaker in the presence of continuous light. *Chlamydomonas* strains used in experiments were *21gr(+)*, *fus1::FUS1-HA(+), hap2(-)*, *hap2::HAP2-HA(-)*, *mar1(-), mar1;hap2(-)*, and *mar1;hap2;MAR1-FLAG(-)*. New strains generated for and used in this study were made by transformation of aforementioned strains and include *mar1::MAR1-FLAG(-), mar1::MAR1-FLAG-MP141-42QQ(-),* and *mar1;hap2::MAR1-FLAG-MP141-142QQ(-)*. Names denote strain origins and genotype; lowercase italics indicates the presence of a mutant gene, uppercase italics indicates the presence of the wild-type gene, double colons (::) indicate strains where transgene introduction was done through direct transformation by electroporation, and semi-colons (;) indicate the presence of a transgene in a progeny strain that was introduced through genetic crosses of a previously transformed parental strain. Further descriptions of the transgenic strains used in this study are in the Key Resources Table.

### *Chlamydomonas* transformation

*Chlamydomonas* cells were transformed by electroporation^86^ using SbfI-linearized plasmid DNA encoding MAR1^21^ with or without the site-directed mutations indicated (Genscript) and a C’terminal 3× FLAG tag. Briefly, mid-log phase vegetative cells were electroporated in the presence of DNA in a 0.4 cm cuvette using BioRad Gene PulserXcell at 25μF and 0.8kV with no shunt resistor. The Genscript sequences of the mutant plasmid MAR1-FLAG DNAs used for transformation were confirmed by sequencing analysis. Positive transformants were selected for further analysis based on their growth on TAP-agar plates containing zeocin (10 μg/mL) and their expression of the FLAG-tagged MAR1 protein in gamete lysates.

## METHOD DETAILS

### Cloning, protein expression and purification of recombinant rMAR1 and rFUS1

Codon*-*optimized DNA sequences encoding the full-length ectodomains of MAR1 (CRR+PRR) and FUS1 (IgL1-7) were synthetized by GenScript and cloned into a modified pMT/BiP plasmid containing a metallothionein-inducible promoter, in-frame with the BiP signal sequence. Shorter ectodomain variants including MAR1 CRR (residues 26-161) and FUS1 IgL 1-4 (residues 17-479) were generated using standard PCR methods. MAR1 constructs featured a C-terminal enterokinase site followed by a double strep-tag, while in FUS1 constructs the strep-tag was replaced by a 6xHis-Tag.

Point mutations in CRR residues 56, 111, 133, 141, 142 and 145 were introduced for pull down assays to test MAR1-FUS1 interaction in solution. To reduce glycosylation heterogeneity in MAR1 CRR, which significantly hindered crystallization of the complex, the N residues at positions 56, 64, 72 and 75 were mutated to Q. These mutations had no impact on protein yield or binding to FUS1 constructs. All mutations were introduced using standard PCR methods.

*Drosophila* S2 cells (ATCC CRL-1963) were cultured at 28°C in serum-free media (HyClone SFM4Insect, Cytiva) supplemented with 50 units/ml of penicillin and 50 units/ml of streptomycin. Cells were co-transfected with different combinations of MAR1 and FUS1 constructs along with the pCoPuro plasmid for puromycin selection of stable transfectants, using the Effectene transfection reagent (Qiagen) according to the manufacturer’s instructions. Cell cultures were further supplemented with 7 μg/ml of puromycin at 3d post-transfection.

For pull-down assays, a FUS1 stable cell line was initially generated with puromycin selection. New cell lines were generated by transfecting this FUS1 cell line with equal amounts of plasmids encoding MAR1 CRR wild-type or the mutants listed above along with pCoBlast plasmid for Blasticidin selection. Selection of stable transfectants was performed by supplementing cell culture medium with 7 μg/ml of Blasticidin at 3d post-transfection.

For large scale protein production, cell-cultures were expanded in 3L spinner flasks until reaching cell densities of ~1×10^7^ cells/ml. At this point, protein expression was induced by activating the metallothionein promoter with 4 μM CdCl2. At five days post-induction, cells were sedimented, and the supernatant was concentrated to 50 ml on a VivaFlow ultrafiltration device (10 kDa MWCO, Sartorius). MAR1- and FUS1-interacting constructs were then purified by affinity chromatography on a StrepTactin column (Cytiva), followed by size exclusion chromatography on a Superdex 200 10 300 column pre-equilibrated with buffer containing 10 mM Tris-HCl pH 8.0 and 100 mM NaCl. Eluted fractions were analyzed by SDS-PAGE under reducing conditions and those fractions containing MAR1 alone or interacting with FUS1 constructs were concentrated and stored at −80°C until use.

### Crystallization and X-ray data collection

The complex between FUS1 IgL1-4 and the MAR1 CRR glycosylation mutant was treated with EndoH glycosidase (New England Biolabs, NEB) for 24h at room temperature followed by enzymatic digestion of the C-terminal tags by incubation with enterokinase (NEB) for an additional 24h at 4°C. The protein solution was subsequently passed through Strep-tactin (Cytiva) and Ni-NTA (Cytiva) columns to retain the uncleaved protein complex. The column flow-through was concentrated and injected in a Superdex 200 10 300 column in the same conditions described above. The central fractions of the chromatographic peak corresponding to the protein complex were pooled and concentrated to 8 mg/ml. Crystals of the complex were obtained by the sitting drop method mixing 0.2 µl of protein solution with an equal volume of the crystallization solution containing 1.2 M Potassium sodium tartrate tetrahydrate, 0.1 M Tris-HCl pH 8.5 and 1 µM Anderson-Evans polyoxotungstate (Jena Bioscience). Hexagonal salt crystals appeared at day 5 and served as nucleation agents for rectangular thin plates than showed up 15 days after. These crystals were fished in crystallization solution supplemented with 15% v/v of glycerol and subsequently flash-frozen in liquid nitrogen. X-ray diffraction data were collected on the Proxima 2 beamline at the SOLEIL Synchrotron (Saint-Aubin, France). The data were reduced using the XDS-package and the structure was solved by molecular replacement with the Phaser Software using predicted models of MAR1 and FUS1 generated with Alphafold 2. The model was manually corrected in COOT and refined with phenix.refine.

### SDS-PAGE and immunoblotting

To make cell lysates, 2.5×10^7^ gametes in 225 μL of N-free medium were lysed by mixture with 75 μL of 4x SDS-PAGE protein sample buffer (160 mM Tris, 4.0% SDS, 20.0% glycerol, 4 mM EDTA, and 0.0006% Bromophenol blue) with reducing agent (TCEP, 5 mM) 5 m at 95°C for detection of MAR1-FLAG proteins, or without reducing agent 5 m at 42°C for detection of HAP2-HA homotrimer (Zhang et al., 2021). 4.2×10^6^ cell equivalents were subjected to electrophoresis on SDS polyacrylamide tris-glycine gels (7%), transferred to PVDF membranes, blocked in 5% milk + TBSt buffer, and probed with either rat anti-HA (1/1000; Sigma, 3F10) or mouse anti-FLAG (1/5000, Sigma, M2) primary antibodies, followed by either goat-anti-rat HRP or goat anti-mouse HRP (1/5000; Millipore) secondary antibodies diluted in 0.5% milk + TBSt buffer. Tubulin was used as a loading control and was detected by probing with mouse anti-α-tubulin primary antibodies (1/5000; Sigma, B512). Immunoblots were developed with SuperSignal West Femto (Thermo Scientific) and scanned with a C-digit blot scanner (LI-COR). Images were processed using Image Studio software (LI-COR) and Adobe Illustrator (V28.5). In some blot images, a white space between lanes indicates cropping of an empty lane.

### Pull-down assays

Protein expression was induced on stable cell lines co-expressing FUS1 IgL1-4 and wild-type or mutated MAR1 CRR. 2 ml of cell culture supernatant was collected and treated with 3 μl of Biolock (IBA Lifesciences) per ml of medium. The supernatant was incubated with MagStrep type 3 XT beads (IBA Lifesciences) for 1h at 4°C with constant agitation. The beads were spun down and washed 3X with buffer containing 10 mM Tris (pH 8) and 100 mM NaCl for 30 m at 4°C. Protein elution was performed using buffer containing 2.5 mM desthiobiotin. The eluted fraction was analyzed by SDS-PAGE and immunoblotting, using HRP-conjugated anti 6xHis-tag mouse antibodies (Proteintech) and anti-strep mouse antibody.

### Protease treatments

Protease treatments to assess cell surface localization of WT and mutated MAR1-FLAG on live gametes were performed as described previously^21,87^ with minor modifications. Briefly, 2.5×10^7^ live *minus* gametes were incubated in the presence and absence of 0.05% pronase (1.0 mg/mL) for 20 m followed by 3 washes in cold N-free media containing 1 mM PMSF, protease inhibitor cocktail (Sigma) and 1% BSA (Sigma), then lysed in SDS-PAGE sample buffer at 95°C for 5 m with 5 mM TCEP prior to SDS-PAGE and immunoblotting.

### Indirect immunofluorescence microscopy

Immunofluorescent staining of gametes was performed as described previously^21^ with minor modifications. Briefly, approximately 1×10^7^ live gametes in N-free medium were allowed to settle and adhere for 10-15 m onto 0.1% poly-L-lysine coated 22×22 mm coverslips. Excess media and non-adhered cells were gently removed by pipetting before adding 100 μl of 4.0% paraformaldehyde for 10 m. Coverslips were then plunged into ice-cold methanol and incubated at −20°C for 20 m, washed 3 × with PBS at 20°C (5 m / wash), washed 2 × with PBST (PBS + 0.5% Triton-X 100), and blocked for 1 h with blocking serum (5% goat serum, 0.1% BSA, 0.5% Triton-X 100, 1% cold-water fish gelatin in PBS). Coverslips were then incubated with anti-FLAG antibodies (1/500 dilution, Sigma, M2) in blocking serum for 2 h at 20°C or overnight at 4°C, washed 3 × with PBSt (PBS + 0.5% Triton-X 100), incubated for 1 h at 20°C in the dark with Alexa Fluor 594-conjugated goat anti-mouse (Invitrogen) secondary antibodies diluted 1/500 in blocking serum, washed 3 × with PBSt and coverslips were mounted using Prolong Gold antifade reagent (Invitrogen). The immunofluorescence described above were used to determine the localization of rMAR1 protein bound to WT(+) and *fus1*(+) gametes with the following modifications: activated, live gametes were incubated with 150 μg/mL of rMAR1 protein for 15 m, before 3 × washing with N-free media and fixation of gametes in 4% paraformaldehyde for 10 m. Fixed gametes were attached to the 0.1% poly-L-lysine coated coverslips. An anti-STREP mouse antibody (1/500) was used to detect the rMAR1 protein in these samples, followed by an Alexa Fluor 488-conjugated goat anti-mouse secondary antibody (1/500, Invitrogen). Immunofluorescence images of gametes were captured on a Leica Stellaris 8 FALCON laser scanning confocal microscope with a Leica 63x/1.4 NA oil objective lens. Images are individual z-sections or z-stack composites as indicated in the figure legends and were processed with Leica LASAF software and Adobe Illustrator (V28.5).

### Phase-contrast and differential interference contrast (DIC) microscopy

Phase-contrast microscopy was used in bioassays to quantify adhesion and fusion and to capture images of quadri-ciliated zygotes and gamete pairs adhered by their mating structures. For phase-contrast imaging, a Zeiss Universal Microscope equipped with a trinocular-mounted CMOS 13″ LCD Display Digital Camera and Zeiss 40x objective lens was used to capture still images from glutaraldehyde-fixed, mixed gamete samples containing an ~1/5 dilution of Lugol’s iodine solution. DIC microscopy was used to assess ciliary adhesion between mt+ and mt− gametes (see Supplemental Fig. S4) with an inverted Axiovert 135 microscope equipped with an WF16X eyepiece adaptor for image and video capture by a mobile device and a 5x/0.15 NA DIC Epiplan NEOFLUAR objective lens.

### Scanning electron microscopy

Gametes were washed 10 × in sterile-filtered N-free media, equal numbers of ectosome-treated HAP2-HA-ALA2 mt− gametes were mixed with mt+ gametes for 30 m, fixed 60 m in Parducz fixative (1 part saturated HgCl_2_, 6 parts 2%OsO_4_)^88^, suspended in fixative, and transferred onto 0.1% poly-L-lysine coated coverslips for an additional 30 m. Coverslips were washed 10 × in ddH2O (2 m/wash), then dehydrated by 3 sequential washes (2 m/wash) in 70%, 95%, and 100% ethanol solutions. Coverslips were critical point dried from CO_2_ and mounted on stubs over double-sided tape before being sputter coated with gold–palladium (60%:40%) (Balzers Med 010) and observed using a scanning electron microscope (Hitachi S-4700 FESEM) at an accelerating voltage of 5 kV in high vacuum.

### Chlamydomonas bioassays

Assays to quantify gamete activation, mating structure membrane adhesion and cell-cell fusion were performed as previously described^21,71^. Briefly, gamete activation was assayed by cell wall loss and quantified by the amount of OD435 detectable chlorophyll released into the supernatant upon treatment of mixed gametes with a detergent buffer^89,90^. Equal numbers of *plus* and *minus* gametes of the indicated genotypes were mixed for 10 m at a concentration of 5 × 10^7^ cells/mL then added to 1.6 volumes of 4°C N-free medium containing 0.075% Triton-X 100 and 5 mM EDTA (pH8), vortexed, centrifuged at 8700 × g for 30 s, and the OD435exp of the supernatant was measured using a Nanodrop 2000 spectrophotometer. The OD435max for individual samples (representing maximal, 100% cell wall loss) was obtained after resuspension and a 12 s sonication of samples with a Microson XL sonicator. Percent gamete activation was calculated as: (OD435exp/ OD435max) × 100. Mating structure adhesion was assayed in mt+ and mt− gametes mixed in equal numbers at a concentration of 5 × 10^7^ cells/mL for 10 m, followed by fixation in an equal volume of 5% glutaraldehyde. Ciliary adhesions were disrupted by pipetting 20× with a 200 μL tip. Glutaraldehyde fixation coupled with the light agitation from pipetting disrupts flagellar adhesions leaving pairs of mt+ and mt− gametes attached only by their mating structures^91^. The percent of gametes attached by their mating structures was calculated as: [(2 × number of pairs) / (2 × number of pairs + number of single gametes)] × 100. Gamete fusion was assayed by the microscopic enumeration of quadri-ciliated cells from equal numbers of mt+ and mt− gametes mixed for 10 m and fixed in an equal volume of 5% glutaraldehyde. The percent of gamete fusion was calculated as: [(2 × number of quadri-ciliated cells) / (2 × number of quadri-ciliated cells + number of single gametes)] × 100. For adhesion and fusion bioassays, fixed samples were stored in 4°C overnight and at least 200 events (quadri-ciliated cells, single cells, or pairs) were counted in each biological replicate.

To assess the effects of rMAR1 on the ability of mt+ gametes to undergo mating structure adhesion and gamete fusion, mt+ gametes (5.0 × 10^7^ cells/mL) were activated by incubation for 1 h in db-cAMP buffer (15 mM db-cAMP, 0.15 mM papaverine, and 40 mM HEPES, pH 7.0^23^), incubated in 150 µM WT or mutant rMAR1 or buffer (10mM Tris and 50mM NaCl) for 20 m, and mixed with equal numbers of mt− gametes for 10 m before fixation in 2.5% glutaraldehyde.

## QUANTIFICATION AND STATISTICAL ANALYSES

### Statistical Analysis

Data were analyzed using Prism 10.2.3 software (GraphPad). Results are shown as box and whisker plots with the box midline representing median, box size representing standard deviation and whiskers showing data minima and maxima. All individual biological replicates for each sample are also overlaid on plots as open circles. Significance was defined as a p-value of <0.05. A non-parametric ANOVA, the Kruskal-Wallis test, and Dunn’s post-test was used to test for significant differences among samples. The sample sizes (n-values) are indicated in the figure legends.

## SUPPLEMENTAL FIGURE LEGENDS

**Figure S1.**
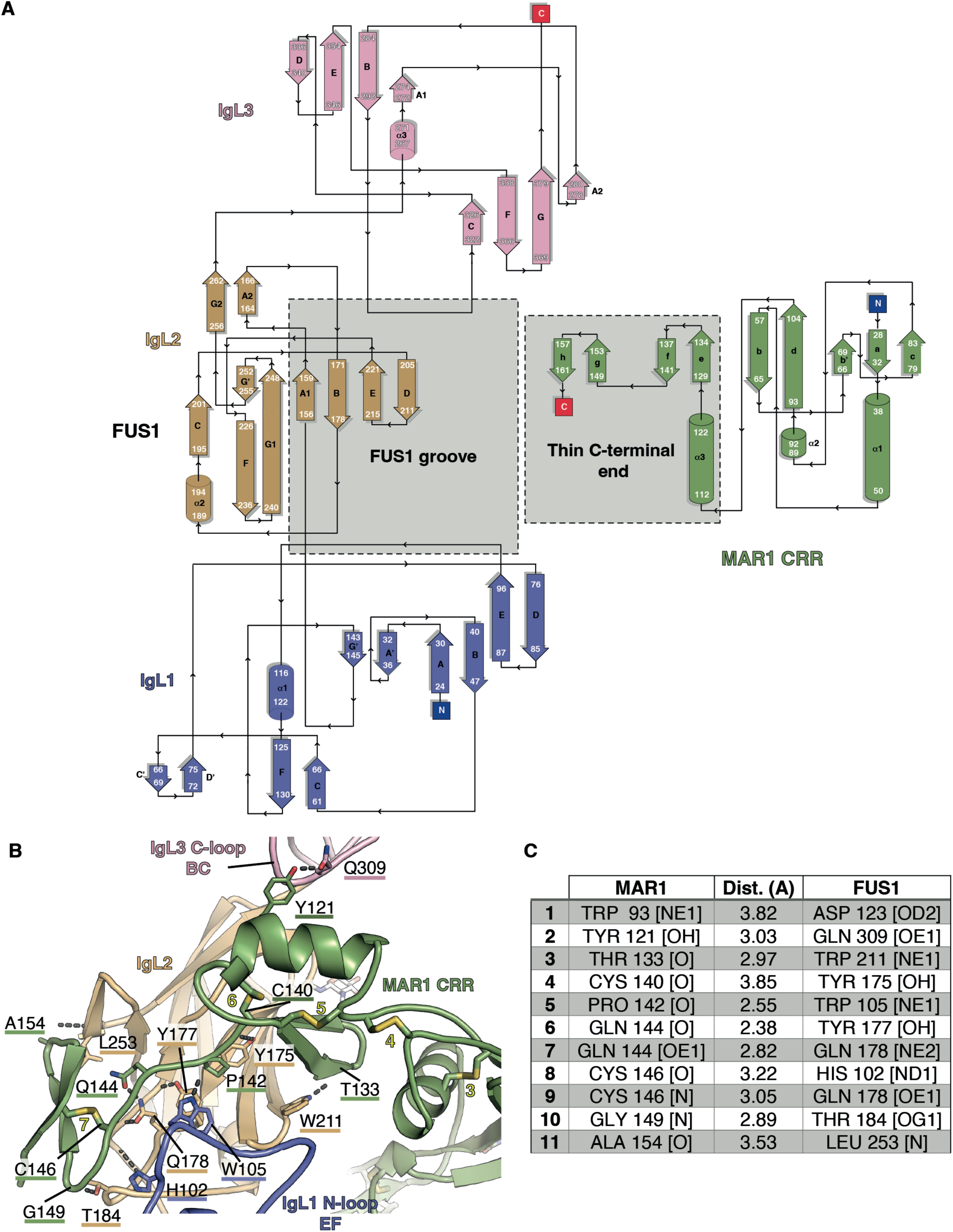
Topology diagram of the complex MAR1 CRR-FUS1 IgL1-3 and description of its main polar interactions. (A) Schematic representation of the interaction MAR1-FUS1 depicting the secondary structure elements (β-strands and helices shown as arrows and cylinders, respectively) of each chain and the regions involved in the interaction between the two proteins (highlighted by gray squares). White labels indicate the amino acid positions at the boundaries of each secondary structure element. (B) Close-up view of the MAR1-FUS1 interaction in ribbon representation. Residues involved in hydrogen bonding between chains are labeled and displayed in stick representation. The interactions are indicated with gray dashed lines. The accompanying table summarizes the residues participating in these interactions and the estimated distance between hydrogen bond acceptor and donor.

**Figure S2.**
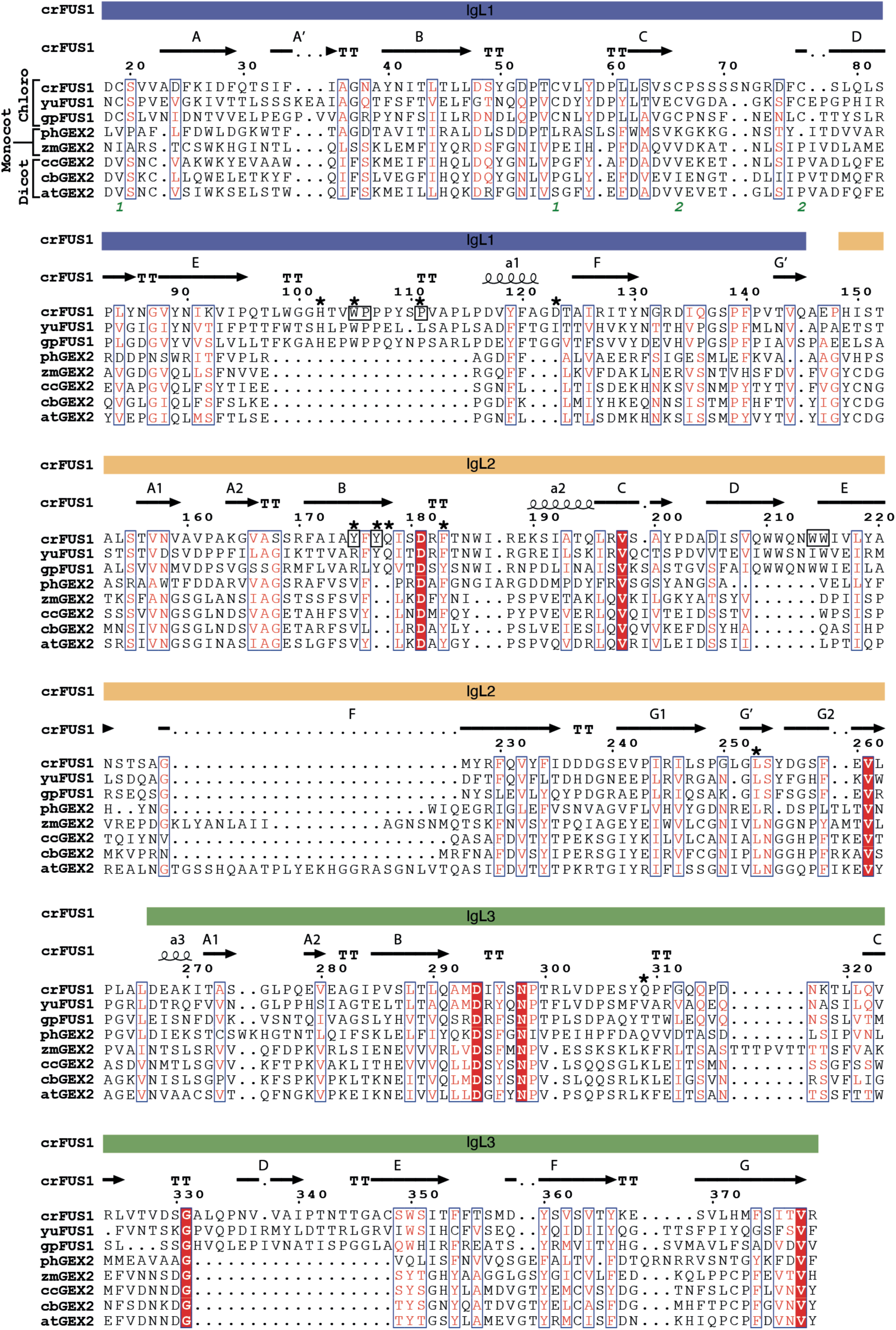
Amino acid sequence alignment of representative FUS1 from Chlorophytes and GEX2 proteins from monocot and dicot plants. The amino acid sequence of *Chlamydomonas reinhardtii* FUS1(crFUS1), *Gonium pectoral* FUS1(gpFUS1), *Yamagishiella unicocca* FUS1 (yuFUS1), *Panicum hallii var. hallii* GEX2 (phGEX2), *Zea mays* GEX2 (zmGEX2), *Citrus clementina* GEX2 (ccGEX2), *Capsicum baccatum* GEX2 (cbGEX2) and *Arabidopsis thaliana* GEX2 (atGEX2) were aligned using the MUSCLE algorithm and rendered with ESPript 3. crFUS1 IgL domain organization and boundaries are indicated by colored bars on top of the alignment. Secondary structure elements are labeled and indicated with black arrows (β-strands) and spirals (helices). crFUS1 residues involved in hydrogen bonds interactions with MAR1 are indicated with black stars. Residues forming the aromatic cage that interacts with MAR1 residues M141 and P142 are enclosed in black frames. Red boxes highlight strict amino acid identity whereas red characters and blue frames indicate similarities in a group or across a group of species, respectively. Green numbers indicate conserved cysteines participating in disulfide bonds in FUS1 proteins.

**Figure S3.**
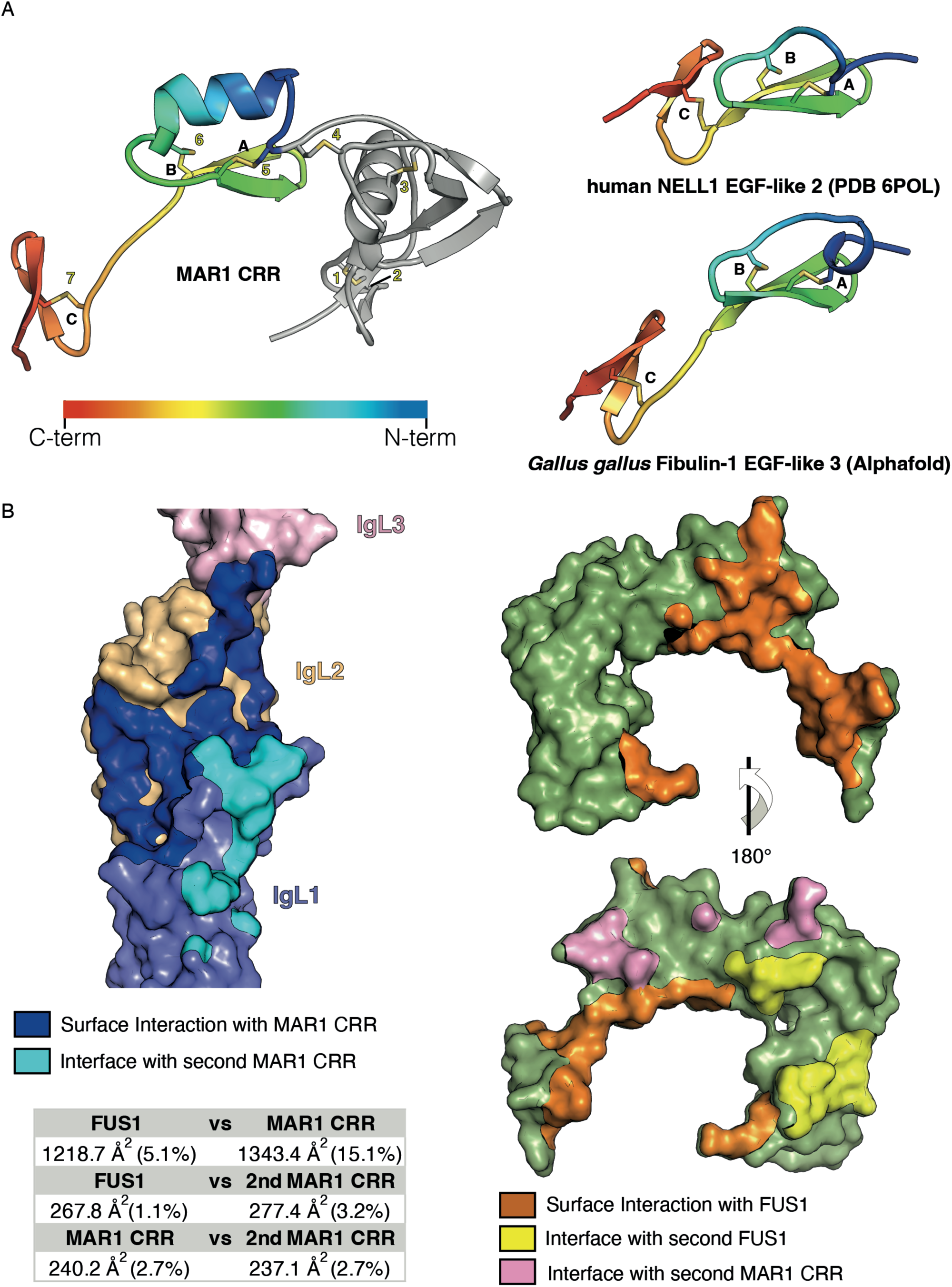
Topology of MAR1 CRR C-terminal end and main interaction surfaces in the complex MAR1-FUS1. (A) Comparison of the secondary structures of MAR1 CRR C-terminal “thin end” (residues 112 to 161) with a representative EGF-like domain from human NELL-1 and a predicted EGF-like from *Gallus gallus* Fibulin-1. EGF-like domains are colored in a rainbow gradient from N-terminus (blue) to C-terminus (red). (B) Main interaction surfaces among different chains in the MAR1 and FUS1 assembly. FUS1 and MAR1 are represented as surfaces and colored by domains similarly as in Figure 1. The interaction surface of one protomer with another chain is highlighted with a distinctive color on each surface. The Buried surface areas (BSA) of the main interaction surfaces in the MAR1-FUS1 assembly were calculated with the PISA algorithm. The percentage of the total protein surface involved in the BSA is indicated in brackets.

**Figure S4.**
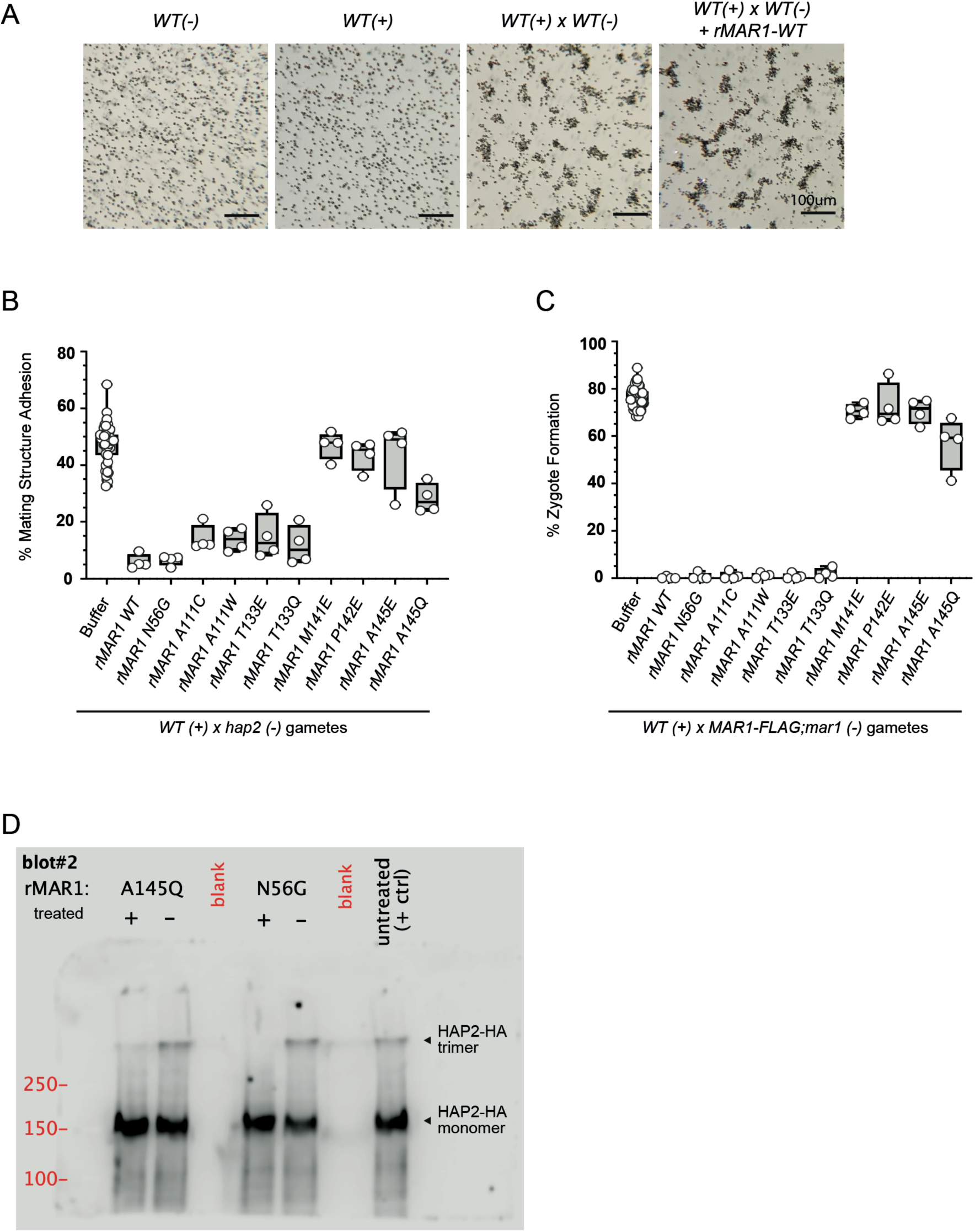
Additional rMAR1 mutant data. For clarity, key treatment conditions for the recombinant MAR1 competition bioassays are highlighted. Multiple rMAR1 point mutants, however, were tested both within and outside the key MAR1-FUS1 interaction region we identified in the co-crystal structure. (A) Gamete activation was unaltered by the presence of wild-type recombinant MAR1 protein in the competition assays. Brightfield microscopic images of wild-type, unmixed mt− and mt+ gametes; wild-type mt+ and mt− gametes without rMAR1 10 min after mixing; and wild-type mt+ and mt− gametes with rMAR1 10 min after mixing (left to right, respectively) showed typical ciliary agglutination. (B-C) Gamete adhesion and fusion are blocked by rMAR1 protein with mutations that are not in the MAR1 interface with FUS1. Activated WT(+) gametes were incubated with rMAR1 WT, single point mutation versions of rMAR1, or a buffer control, and an equal number of activated, fusion defective, *hap2(-)* gametes (B, adhesion); or fusion-competent, *MAR1-FLAG*;*mar1(-)* gametes (C, fusion) were added, and the percentages of cells in pairs (B), or as quadri-ciliated zygotes (C), were assessed after 10 m. (D) Similar to WT rMAR1 incubations shown in Fig.3G, HAP2 trimer formation was blocked by pre-incubation of mt+ gametes with rMAR1 containing single point mutations outside of the MAR1 interface with FUS1. Activated WT(+) gametes were treated with rMAR1 A145Q, N56G, or a buffer control. Then an equal number of activated, *fh1(-)* gametes was added for 10 m and the cell mixture was lysed in non-reducing sample buffer at 42°C and subjected to immunoblotting to assess HAP2-HA trimer formation.

